# Neural geometrodynamics, complexity, and plasticity: a psychedelics perspective

**DOI:** 10.1101/2023.08.14.553258

**Authors:** G. Ruffini, E. Lopez-Sola, J. Vohryzek, R. Sanchez-Todo

## Abstract

We explore the intersection of neural dynamics and the effects of psychedelics in light of distinct timescales in a framework integrating concepts from dynamics, complexity, and plasticity. We call this framework *neural geometrodynamics* for its parallels with general relativity’s description of the interplay of spacetime and matter. The geometry of trajectories within the dynamical landscape of “fast time” dynamics are shaped by the structure of a differential equation and its connectivity parameters, which themselves evolve over “slow time” driven by state-dependent and state-independent plasticity mechanisms. Finally, the adjustment of plasticity processes (metaplasticity) takes place in an “ultraslow” time scale. Psychedelics flatten the neural landscape, leading to heightened entropy and complexity of neural dynamics, as observed in neuroimaging and modeling studies linking increases in complexity with a disruption of functional integration. We highlight the relationship between criticality, the complexity of fast neural dynamics, and synaptic plasticity. Pathological, rigid, or “canalized” neural dynamics result in an ultrastable confined repertoire, allowing slower plastic changes to consolidate them further. However, under the influence of psychedelics, the destabilizing emergence of complex dynamics leads to a more fluid and adaptable neural state in a process that is amplified by the plasticity-enhancing effects of psychedelics. This shift manifests as an acute systemic increase of disorder and a possibly longer-lasting increase in complexity affecting both short-term dynamics and long-term plastic processes. Our framework offers a holistic perspective of the acute effects of these substances and their potential long-term impacts on neural structure and function.

## 1 Introduction

In this paper, we explore new perspectives to interpret changes in the brain’s landscape and connectivity, focusing on the subtle interplay between structural and dynamical aspects across timescales (fast, slow, and ultraslow). Our primary goal is to present a framework that enhances the understanding of the intricate relationships among brain dynamics, complexity, structure, and plasticity. This framework, which we call “neural geometrodynamics”, draws on principles from non-linear dynamics and is further inspired by conceptual links to general relativity in physics.

In describing neural dynamics, we will refer to the mathematical formalism of neural mass models (NMMs), although other computational neuroscience formulations are equally relevant [1, 2]. Neural mass models have been extensively utilized to model various brain activities, from localized brain functions to the coordinated activity observed in different brain regions. By employing mathematical formulations that include essential features like synaptic connectivity and neuronal excitability, NMMs enable the simulation and analysis of complex brain activities in various dynamic regimes [3]. NMMs are particularly useful because they provide a link between the mesoscopic physiological scale and macroscopic brain function, allowing for the connection of effects on neurons at the molecular level, such as those of psychedelics, with those of whole-brain connectivity [4, 5].

Analyzing the effects of psychoactive neuroplastogens (psychedelics such as psilocybin or LSD) serves as an illustrative case of the framework, given the immediate and potentially lasting plastic changes these substances can provoke in the brain [6]. By altering neural dynamics and connectivity, psychedelics are thought to induce both transient and sustained shifts in cognition and perception [7]. Several studies underscore the role of psychedelics in inducing neuroplasticity with antidepressant effects, revealing mechanisms at molecular, synaptic, and dendritic levels [8, 9], and with significant potential for treating neuropsychiatric disorders [10, 11], although the duration and permanence of these effects remain to be fully understood.

Recent conceptual perspectives have enhanced our understanding of the brain’s response to psychedelics, combining biological, dynamical systems, complexity science, and artificial intelligence viewpoints. The REBUS (RElaxed Beliefs Under pSychedelics) framework [12], grounded in the Free Energy Principle (FEP) and the entropic and anarchic brain models, offers a perspective on the effects of psychedelics on the brain whereby psychedelic action results in the collapse of brain functional hierarchies or, in other words, in the “flattening of the landscape” of brain’s dynamics to allow the brain state to escape a deep local minimum. The term *annealing* is also used in this context in relation to physical annealing in metallurgy and simulated annealing in numerical optimization [13].

Consequently, it has been argued that the observed expansion of the repertoire of functional patterns elicited by hallucinogenic substances can be associated with an increase in entropy in brain dynamics [14, 15], with the brain moving to a more disordered state from a relaxation of high-level cognitive priors [12, 16]. This may lead to a favorable context for conducting psychotherapy [12, 17, 18]. Studies on functional neuroimaging regarding psilocybin and LSD effects have shown initial evidence of the mechanistic alterations on brain dynamics at the network level, with the majority of the findings suggesting a relative weakening of usual functional configurations giving place to an increase of brain entropy, global functional integration, and more flexible brain dynamics [14, 19–28]. As mentioned above, these changes are traditionally reflected in the complexity of neural dynamics, which can be evaluated using various techniques such as criticality measures [29, 30], complexity measures [31], connectome harmonic decomposition [23–25], control theory [26] and Ising (or spinglass) modeling [32, 33].

For example, Ising modeling of psychedelics has shown that the increased complexity of brain dynamics under LSD (e.g., increased Ising temperature, Lempel-Ziv, and the Block Decomposition Method complexity) is associated with a decrease of interhemispheric connectivity — especially homotopic links [34], corroborating earlier modeling studies suggesting the central role of long-range connections in controlling phase transitions [35].

The observed push of brain dynamics towards disorder and away from criticality aligns with the REBUS and FEP frameworks, which link the vividness of experience to the entropy of brain activity. At the same time, the notion that a wakeful brain exhibits dimensionality reduction and criticality features that are disrupted by the effect of psychedelics is also predicted by an algorithmic perspective on consciousness [16, 36, 37], where the psychedelic shift towards disorder is associated with a disruption of the world-modeling/world-tracking circuits in the brain.

Another feature of brain dynamics related to the collapse of higher-order cognitive functions under psychedelics in the REBUS framework is the hierarchical organization along the uni-to trans-modal functional gradient [38]. This asymmetry in neural activity reflects the bottom-up and top-down information flows in cognitive processing [39, 40]. This has been suggested to be intimately linked to non-equilibrium dynamics in thermodynamic-inspired frameworks where the level of hierarchy is related to the amount of brain signal irreversibility as well as entropy production [41–43]. Indeed it has been demonstrated that the principal functional gradient collapses under the influence of various psychedelics [44–46].

A related perspective for this paper is the CANAL framework [11] for describing the pathological plasticity of “being stuck in a rut” in certain mood disorders and the potential therapeutic role of psychedelics through the concept of metaplasticity. In contrast to psychedelics, these changes are reflected in neural dynamics with brain signatures of excessively rigid and highly ordered functional states [47]. The CANAL framework has been further extended by establishing connections with deep artificial neural networks (Deep CANAL [48]) to introduce a distinction between two distinct pathological phenomena — one related to fast brain dynamics and their slow and ultraslow counterparts. These distinctions will be naturally integrated into the presented framework (see the Appendix for a figure relating the concepts in the different frameworks).

While the discussion is centered on the effects of psychedelics, the framework proposed here extends more generally to other phenomena related to plasticity, including neurodevelopment, pathological plasticity in mood disorders [49], and interventions that alter brain dynamics like transcranial brain stimulation (tES) [50], transcranial magnetic stimulation (TMS), or electroconvulsive therapy (ECT).

In what follows, we formalize the notions of brain dynamics, plasticity, and their associated timescales and subsequently use them to study the impact of psychedelics on the brain. In the last section, we draw connections between the framework and concepts from general relativity in physics. We hope these parallels will illuminate the complex relationship between the structure and function of brain dynamics. Figure 1 illustrates the reciprocal dynamics between brain states and connectivity as conceptualized in the neural geometrodynamics framework.

**Figure 1:**
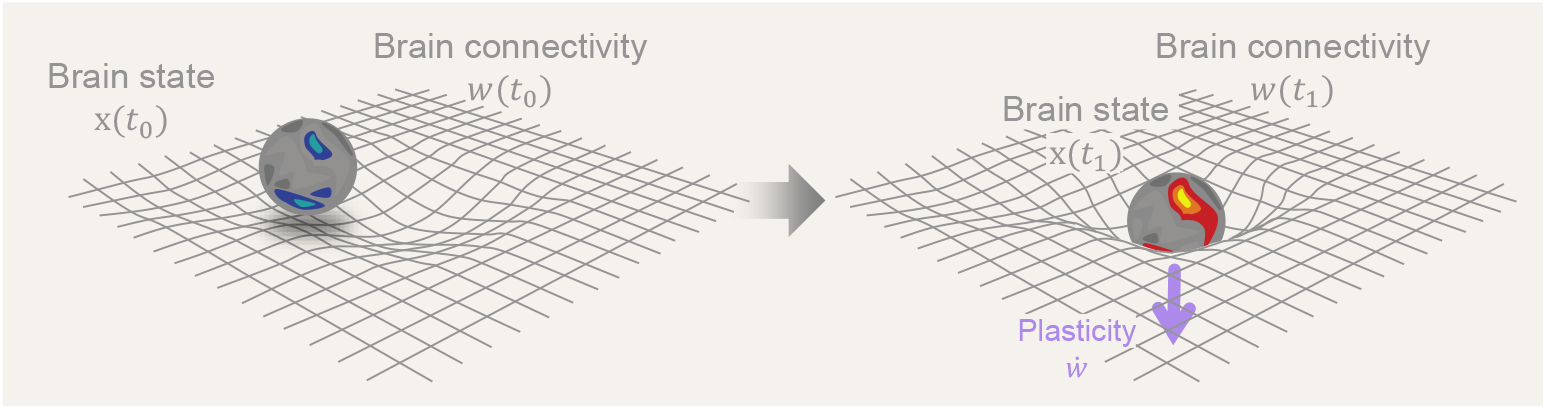
Neural Geometrodynamics: a dynamic interplay between brain states and connectivity. A central element in the discussion is the dynamic interplay between brain state (*x*) and connectivity (*w*), where the dynamics of brain states is driven by neural connectivity while, simultaneously, state dynamics influence and reshape connectivity through neural plasticity mechanisms. The central arrow represents the passage of time and the effects of external forcing (from, e.g., drugs, brain stimulation, or sensory inputs), with plastic effects that alter connectivity (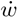, with the overdot standing for the time derivative).

## 2 Dynamics across timescales

The state of a system can be defined by a set of coordinates in phase space: a multidimensional manifold in which each dimension corresponds to one of the variables. For a single particle moving in one dimension, the phase space is two-dimensional, with one axis representing its position and the other representing its momentum. For example, Figure 2 illustrates the phase space of a pendulum with friction. In phase space, and perhaps after some transient period, the possible trajectories of the states of the system lie in a reduced or invariant manifold (an attractor, see Box 1 for a glossary of terms), which we may refer to as the “geometry” or latent “structure” of the phase space. Together, the structure (geometry and topology) of the phase space with its invariant properties can be referred to as the dynamical landscape, where the depth or shallowness of the “valleys” can, in some cases, be interpreted as the stability of the state in that location given some stochastic forcing. For example, in mechanics, the landscape can be labeled by potential energy isolines, e.g., in a physical system such as in the pendulum example in Figure 2 (bottom right), or their generalization, Lyapunov functions [55].

**Figure 2:**
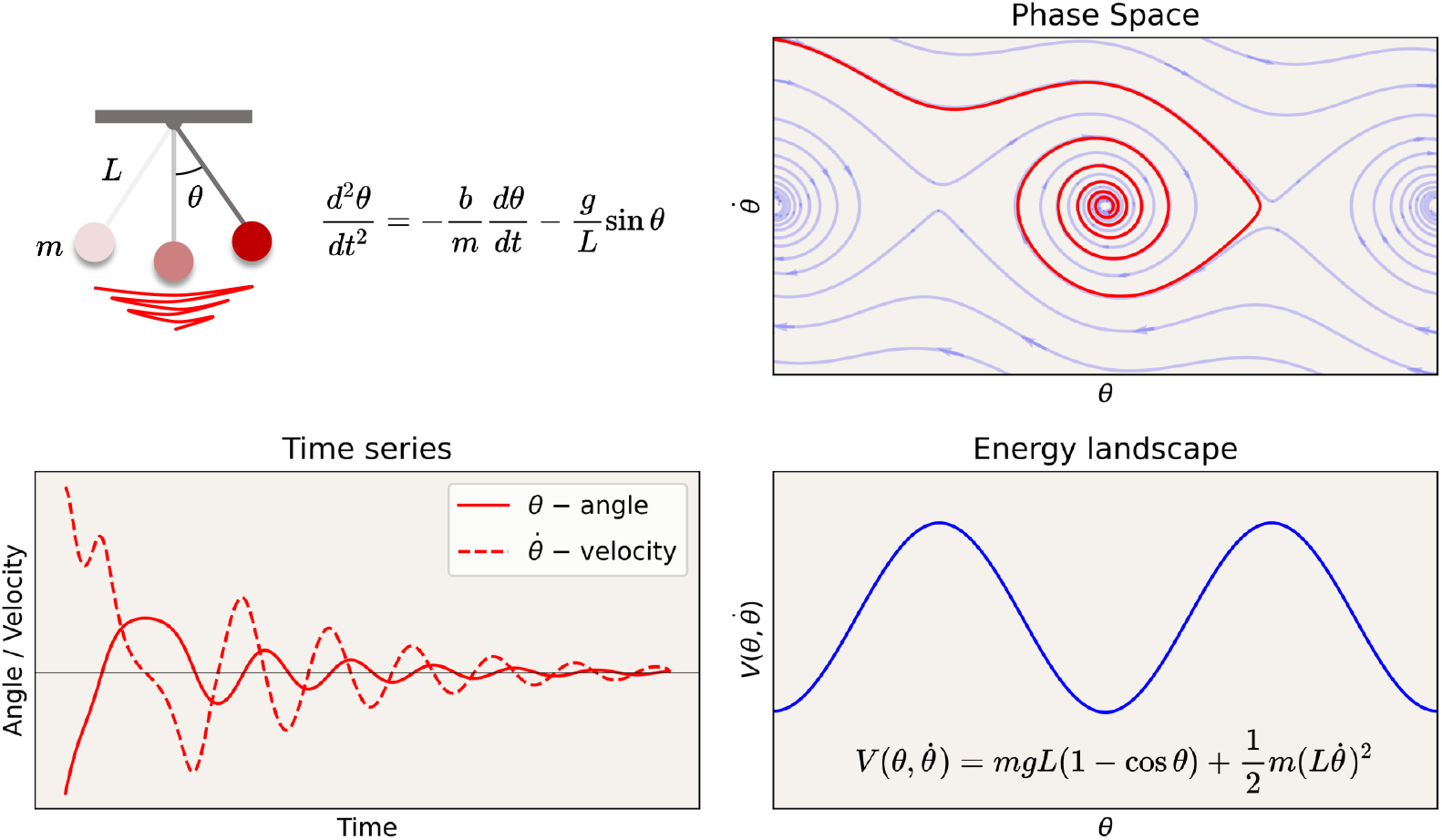
Dynamics of a pendulum with friction. Time series, phase space, and energy landscape. Attractors in phase space are sets to which the system evolves after a long enough time. In the case of the pendulum with friction, it is a point in the valley in the “energy” landscape (more generally, defined by the level sets of a Lyapunov function).

### Fast time: neural dynamics

Here, we discuss the first equation in neural geometrodynamics in the context of neural mass models, but the ideas are applicable more extensively in computational neuroscience. The standard equation we use in neural mass modeling is a multidimensional

#### Box 1

GLOSSARY

##### State of the system

Depending on the context, the state of the system is defined by the coordinates *x* (Eq. 1, fast time view) or by the full set of dynamical variables (*x, w, θ*) – see Eqs. 1, 2 and 3.

##### Entropy

Statistical mechanics: the number of microscopic states corresponding to a given macroscopic state (after coarse-graining), i.e., the information required to specify a specific microstate in the macrostate. Information theory: a property of a probability distribution function quantifying the uncertainty or unpredictability of a system.

##### Complexity

A multifaceted term associated with systems that exhibit rich, varied behavior and entropy. In algorithmic complexity, this is defined as the length of the shortest program capable of generating a dataset (Kolmogorov complexity). Characteristics of complex systems include nonlinearity, emergence, self-organization, and adaptability.

##### Critical point

Dynamics: parameter space point where a qualitative change in behavior occurs (*bifurcation point*, e.g., stability of equilibria, emergence of oscillations, or shift from order to chaos). Statistical mechanics: phase transition where the system exhibits changes in macroscopic properties at certain critical parameters (e.g., temperature), exhibiting scale-invariant behavior and critical phenomena like diverging correlation lengths and susceptibilities. These notions may interconnect, with bifurcation points in large systems leading to phase transitions.

##### Temperature

In the context of Ising or spinglass models, it represents a parameter controlling the degree of randomness or disorder in the system. It is analogous to thermodynamic temperature and influences the probability of spin configurations. Higher temperatures typically correspond to increased disorder and higher entropy states, facilitating transitions between different spin states.

##### Effective connectivity (or connectivity for short)

In our high-level formulation, this is symbolized by *w*. It represents the connectivity relevant to state dynamics. It is affected by multiple elements, including the structural connectome, the number of synapses per fiber in the connectome, and the synaptic state (which may be affected by neuromodulatory signals or drugs).

##### Plasticity

The ability of the system to change its effective connectivity (*w*), which may vary over time.

##### Metaplasticity

The ability of the system to change its plasticity over time (dynamics of plasticity).

##### State or Activity-dependent plasticity

Mechanism for changing the connectivity (*w*) as a function of the state (fast) dynamics and other parameters (*α*). See Eq. 2.

##### State or Activity-independent plasticity

Mechanism for changing the connectivity (*w*) independently of state dynamics, as a function of some parameters (*γ*). See Eq. 2.

##### Connectodynamics

Equations governing the dynamics of *w* in slow or ultraslow time.

##### Fast time

Timescale associated to state dynamics pertaining to *x*.

##### Slow time

Timescale associated to connectivity dynamics pertaining to *w*.

##### Ultraslow time

Timescale associated to plasticity dynamics pertaining to *θ* = (*α, γ*) — v. Eq. 3.

##### Phase space

Mathematical space, also called **state space**, where each point represents a possible state of a system, characterized by its coordinates or variables.

##### Geometry and topology of reduced phase space

State trajectories lie in a submanifold of phase space (the reduced or invariant manifold). We call the geometry of this submanifold and its topology the “structure of phase space” or “geometry of dynamical landscape”.

##### Topology

The study of properties of spaces that remain unchanged under continuous deformation, like stretching or bending, without tearing or gluing. It’s about the ‘shape’ of space in a very broad sense. In contrast, geometry deals with the precise properties of shapes and spaces, like distances, angles, and sizes. While geometry measures and compares exact dimensions, topology is concerned with the fundamental aspects of connectivity and continuity.

##### Invariant manifold

A submanifold within (embedded into) the phase space that remains preserved or invariant under the dynamics of a system. That is, points within it can move but are constrained to the manifold. Includes stable, unstable, and other invariant manifolds.

##### Stable manifold or attractor

A type of invariant manifold defined as a subset of the phase space to which trajectories of a dynamical system converge or tend to approach over time.

##### Unstable Manifold or Repellor

A type of invariant manifold defined as a subset of the phase space from which trajectories diverge over time.

##### Latent space

A compressed, reduced-dimensional data representation (see Box 2).

##### Topological tipping point

A sharp transition in the topology of attractors due to changes in system inputs or parameters.

ODE of the form

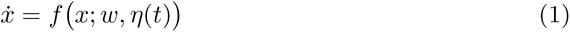

with *x* ∈ ℝ^*n*^ and where *w* denotes connectivity parameters^1^ and where, as usual, a dot over a variable denotes its time derivative. This equation governs dynamics at short time scales (seconds or less) when connectivity parameters *w* are assumed to be constant.

#### Box 2

The manifold hypothesis and latent spaces e manifold hypothesis and latent spaces

The dimension of the phase (or state) space is determined by the number of independent variables required to specify the complete state of the system and the future evolution of the system. The **Manifold hypothesis** posits that high-dimensional data, such as neuroimaging data, can be compressed into a reduced number of parameters due to the presence of a low-dimensional invariant manifold within the high-dimensional phase space [51, 52]. **Invariant manifolds** can take various forms, such as **stable manifolds or attractors** and unstable manifolds. In attractors, small perturbations or deviations from the manifold are typically damped out, and trajectories converge towards it. They can be thought of as lower-dimensional submanifolds within the phase space that capture the system’s long-term behavior or steady state. Such attractors are sometimes loosely referred to as the **“latent space”** of the dynamical system, although the term is also used in other related ways. In the related context of deep learning with variational autoencoders, latent space is the compressive projection or embedding of the original high-dimensional data or some data derivatives (e.g., functional connectivity [53, 54]) into a lower-dimensional space. This mapping, which exploits the underlying invariant manifold structure, can help reveal patterns, similarities, or relationships that may be obscured or difficult to discern in the original high-dimensional space. If the latent space is designed to capture the full dynamics of the data (i.e., is constructed directly from time series) across different states and topological tipping points, it can be interpreted as a representation of the invariant manifolds underlying system.

The external input term *η*(*t*) makes the equations non-autonomous (an autonomous ODE does not explicitly depend on time). This term can refer to external forces providing random kicks to the trajectory or to a more steady and purposeful forcing from unspecified internal systems, external inputs from sensory systems, or external electric fields, for example.

We may think of this equation describing phenomena in **fast time** scales as providing the “structure” for the dynamics of neuronal population state. The fast timescale is set by synaptic transmission (milliseconds) and by ephaptic coupling (electromagnetic waves) [57–60] in a nanosecond or subnanosecond scale [59].

Equation 1 characterizes the **dynamical landscape**, which is established through the geometric structure of the phase space, where trajectories are shaped by the given set of ordinary differential equations. The landscape is determined by the functional form of *f* (*x*; *w, η*(*t*)) and by the parameters *w*, and is analogous to the Neural Activation Landscape proposed in [48]. More specifically, we talk about the landscape as defined by the manifold generated by the motion of trajectories with coordinates *x* ∈ ℝ^*n*^. Typically, trajectories lie in a reduced manifold of dimensionality lower than ℝ^*n*^. The fact that such a reduced space exists means that it can be generated by coordinates in a reduced latent space. The geometry and topology of this reduced space in different states provide a synthetic description of the dynamics and are of special interest [37].

### Slow time: connectodynamics

The landscape, like that on planet Earth, may appear to be static, but in reality, it is not fixed. It also flows in *slow time*. We thus consider changes in connectivity in the system, that is, now *w* = *w*(*t*). We call the potential for such changes *plasticity of the system*. The general form of this equation is 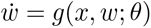, with *θ* standing for a set of parameters controlling plasticity.

To be more concrete, we can think of two types of process: one that modifies the connectivity parameters independently of the system’s state (*ψ*) and another that is a function of the state (e.g., Hebbian plasticity [61], *h*). We express this by writing

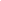

(with the second term understood as not separable into parts where any part is a function of only *w*). This decomposition separates out *state-dependent* (via the term *h*(*x, w*; *α*)) and *state-independent plasticity* (with *ψ* (*w*; *γ*)) processes. The set of parameters *θ* is similarly decomposed as *θ* = (*α, γ*): we separate out the plasticity-controlling parameters in order to differentiate the *state-dependent (α)* and *state-independent (γ)* plasticity control parameters (e.g., Hebbian vs. drug-enhanced structural plasticity [11]). The parameters (*α, γ*) may vary in time to reflect, for example, the effects of drugs. The dynamics of these parameters are formalized in the next section.

Hebbian plasticity is the most prominent example of state-dependent plasticity [61]. State dependence implies that state-related concepts such as system temperature, phase transitions, and critical phenomena are relevant for the study of the dynamics of plasticity. In particular, within the scope of slower “slow time” (taking place over many hours), we include *homeostatic plasticity* [62–65], which may itself target desired complexity states as a homeostatic goal [66, 67]. In the case of state-independent plasticity, there are numerous candidates for these plastic processes, such as heterosynaptic plasticity [68] or critical-period plasticity [69].

In summary, the functions *h* and *ψ* with parameters *α* and *γ* regulate *connectodynamics*, defining where and how fast the effective connectivity will change in a state-dependent or state-independent way.

These connectodynamics differential equations define a new dynamical landscape, which we can call the *plasticity landscape* (analogous to the Synaptic Weight Landscape in [48]). The state *w* in this plasticity landscape will determine the shape of the neural dynamics landscape.

### Ultraslow time: metaplasticity

Plasticity is required to adapt to a changing environment [70], and the environment may change at different rates at different times. Plasticity in the healthy brain should match this variation in the character of dynamics accordingly. This is analogous to the situation in biology, where optimal mutation rates ensure successful adaptation in a tradeoff with genetic integrity [71]. More specifically, the plasticity-regulating parameters *α* and *γ* in Equation 2 should adapt to changes in the environmental conditions.

In pathological cases, plasticity levels can either become overly exuberant, reflecting the notion of catastrophic forgetting in artificial neural networks, or impoverished and rigid, reflecting general plasticity loss [48]. These scenarios can be tentatively related to certain neurological and psychiatric conditions. For example, reduced plasticity could underlie conditions such as major depressive disorder, obsessive-compulsive disorder, anxiety, or substance abuse [48, 72].

To account for the dynamics of plasticity, we allow the plasticity parameters to be dynamic, i.e.,

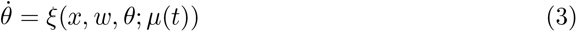

This equation is again state-dependent, allowing the system to respond to changes in the neural dynamics (with state dynamics as drivers of plasticity parameter regulation [73]), including critical phenomena (changes in criticality regime [66]) and complexity. Plasticity dynamics reflect changes in the parameters regulating state-dependent (Hebbian) plasticity (changes in *α*) during neurodevelopment, and state-independent plasticity, such as the ones induced by psychedelics in the acute or post-acute phases (changes in *γ*). Finally, this equation is a function of other parameters and non-autonomous terms (*μ* (*t*)), reflecting external perturbations of the system, such as those from drugs. We provide analogies in the context of sailing and electrodynamics in the appendix to further clarify these concepts.

The dynamics of plasticity presented above reflect a physiological principle well described by Abraham et al. in the definition of metaplasticity [74]

> **Metaplasticity** [ …*] is manifested as* **a change in the ability to induce subsequent synaptic plasticity**, *such as long-term potentiation or depression. Thus, metaplasticity is a higher-order form of synaptic plasticity [74]*.

Thus, metaplasticity and its counterparts are terms used in neuroscience to refer to the plasticity of synaptic plasticity. That is, the idea that the ability of synapses to strengthen or weaken in response to increases or decreases in their activity (which is called synaptic plasticity) can be modulated based on the history of the synaptic activity or other factors (e.g., age, neuromodulatory systems, drugs, or lifestyle [75]). Metaplasticity has important implications for the learning and memory of an organism, as it can regulate the ability of synaptic plasticity to change and adapt over time as required by its environmental context.

We call the set of equations 1,2 and 3 — somewhat whimsically — the equations for *neural geometrodynamics* in reference to the equations of general relativity in physics. We recall that general relativity provides equations defining the dynamics of spacetime geometry (via the “metric”) coupled with matter [76]. Section 4 elaborates further on this parallel.

## 3 Dynamics under psychedelics

Psychedelics like psilocybin and LSD act as agonists or partial agonists for serotonin 5-hydroxytryptamine 2A (5-HT_2A_) receptors, specifically targeting Layer V cortical pyramidal neurons [11, 14, 56, 77, 78], leading to increased neuronal excitability through an increase in excitatory postsynaptic currents and discharge rates in pyramidal neurons [12]. The highest expression of 5-HT2ARs is found on the apical dendrites of Layer 5 pyramidal cells in both cortical and subcortical structures [12, 79]. In the cortex, 5-HT_2A_ receptors are strongly expressed along a steep anteroposterior gradient [80]. When psychedelics bind to these receptors, they can lead to a gradual increase of the excitability of these pyramidal neurons — depolarizing them and making them more susceptible to excitatory inputs such as those associated with glutamate receptors [80] — much as the gain knob in an amplifier. This increased excitability and susceptibility to inputs can lead to changes in the firing patterns of these neurons and alterations in the overall neural circuit activity. Recognized for their potent and immediate impact on the brain, these drugs cause a swift reconfiguration of neural dynamics. As we explain, this immediate effect is represented in our model by state-independent alterations in the connectivity parameters (*w*) (see Figure 3).

**Figure 3:**
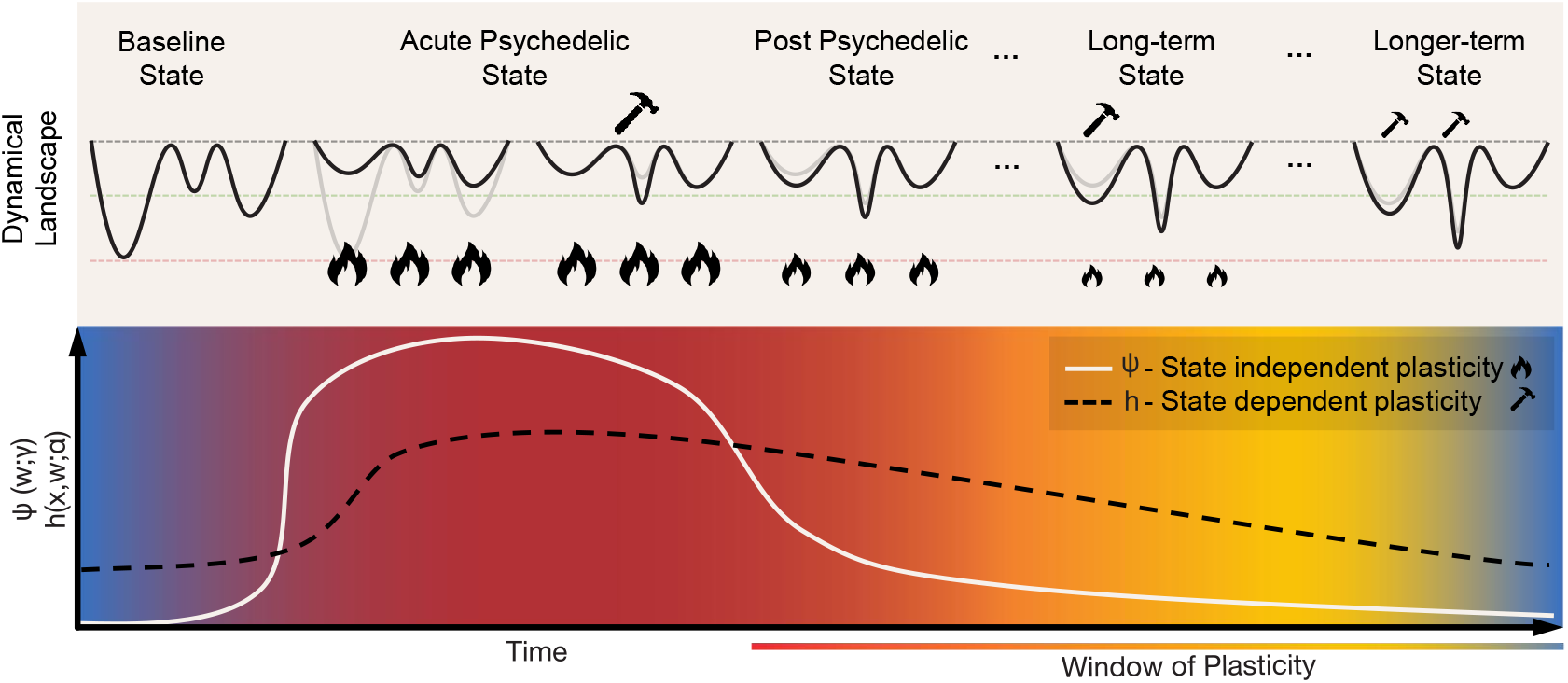
Geometrodynamics of the acute and post-acute plastic effects of psychedelics. The **acute** plastic effects can be represented by rapid state-independent changes in connectivity parameters, i.e., the term *ψ* (*w*; *γ*) in Eq. 3. This will result in the flattening or de-weighting of the dynamical landscape. Such flattening allows for the exploration of a wider range of states, eventually creating new minima through state-dependent plasticity, represented by the term *h*(*x, w*; *α*) in Eq. 3. As the psychedelic action fades out, the landscape gradually transitions towards its initial state but with lasting changes due to the creation of new attractors during the acute state. The **post-acute** plastic effects can be described as a “window of enhanced plasticity”. These transitions are brought about by changes of the parameters *γ* and *α*, each controlling the behavior of state-independent and state-dependent plasticity, respectively. In this post-acute phase, the landscape is more malleable to internal and external influences.

How are these effects represented in Equations 1 and 2? If we include the neuromodulatory nodes in our model — the dorsal raphe and median raphe nuclei in the brainstem are the source of most serotonergic neurons projecting throughout the brain [80] —, the modulation of serotonin receptors could be represented by changes in neuromodulatory connectivity (the subset of *w* parameters in the model connecting the raphe nuclei to other nodes). Alternatively, if neuromodulatory nodes are not explicitly included in the model, for the purposes at hand, we can think of the changes in the excitability of the nodes affected by neuromodulatory inputs as leading to changes in their effective connectivity (*w*) to other nodes (e.g., through an increase of the connectivity of glutamatergic synapses into Layer 5 pyramidal cells).

The abrupt shift induced by psychedelics can be thought of as a transformation of the phase space’s geometry, allowing the neural state to explore new trajectories. This process manifests in an increase of complexity and disorder, which can be measured using various tools in different modalities (e.g., EEG or fMRI BOLD with measures such as entropy, fractal dimension, algorithmic complexity, etc. [29, 31, 34, 81]). The decrease in effective connectivity under LSD (especially in interhemispheric homotopic connections), as inferred using Ising modeling of BOLD signals measured using fMRI imaging, is associated with a subsequent increase in algorithmic complexity [34].

Psychedelic-induced changes in connectivity correspond to a flattening of the dynamical landscape [12] or a destabilization of it [48]. In our framework, the alteration of effective connective results in an immediate and state-independent remodeling of the dynamical landscape during the acute phase of psychedelics, which is represented by the term *ψ* (*w*; *γ*) in the connectodynamics equation (Eq. 2)^2^.

The instantaneous modification of the landscape is, however, ephemeral, gradually fading as the acute effects of the psychedelics wear off. The system returns to near its original geometrical configuration but with lasting influences brought about by the plastic changes resulting from the exploration of new trajectories in the acute phase. These residual changes are captured by the state-dependent plasticity term, *h*(*x, w*; *α*), which reflects changes in connectivity due to Hebbian plasticity that arise from the co-activation of neurons during the psychedelic acute stage.

In the literature, there is an increasing body of evidence suggesting a post-acute phase following psychedelic exposure characterized by a period of enhanced plasticity [11, 12, 82–84]. This phase can be interpreted as an extended window of malleability of the landscape, which could have profound implications for learning and therapy. Such window of plasticity has been related to increased neurogenesis and upregulation of Brain-Derived Neurotropic Factor (BDNF) in humans and mice [8]. The activity-dependent release of BDNF plays a crucial role in selectively strengthening active synapses while weakening inactive ones, a critical process for Hebbian-type plasticity. Intriguingly, recent studies with mice have found psychedelic-induced changes in plasticity and antidepressant-like behavior dependent on the increase of endogenous BDNF and TrkB binding (the receptor of BDNF), but independent from the activation of 5-HT_2A_ [9, 10].

In terms of our model, these two pathways correspond to changes of connectivity through Equation 2 due to a temporary modulation of the parameters *γ* and *α* (i.e., metaplasticity, see Equation 3) upregulating state-independent and state-dependent plasticity processes, respectively. The strong acute-phase increase of state-independent plasticity (*ψ*) would be directly associated with the activation of serotonergic receptors, as discussed above, with a possible gradual decrease during the post-acute phase (solid white line in Figure 3). The sustained increase of state-dependent plasticity (*h*) in the post-acute phase (dashed black line in Figure 3) would be linked to dendritic growth, neurogenesis, upregulation of BDNF, and other related changes. This means that in the post-acute period, the landscape would be more responsive to state changes (itself influenced by external factors), offering a potential mechanism for the long-lasting changes reported after psychedelic experiences. Such external influences are modeled by the external input term *η*(*t*) in the state equation (Eq. 1) and can represent environmental/sensory inputs, psychotherapy, or neuromodulatory brain stimulation techniques such as transcranial electrical current stimulation (tES).

### Dynamics of psychedelics and psychopathology

Recently, psychedelic medicine has emerged as a promising direction for treating mental disorders such as depression or addiction [85]. The nuanced interaction between the brain’s neurophysiology and the emergent brain activity underlies the pathophysiology of mood disorders, often resulting in a persistent and maladaptive rigidity in cognitive and emotional processes [86]. Such changes to the brain’s neurophysiology can be explained through the CANAL framework whereby pathological plasticity, often caused by a traumatic event, asserts itself and dominates brain activity, driving the brain state to be “stuck in a rut” [11], i.e., a deepening minimum in the dynamical landscape (see Figure 4).

**Figure 4:**
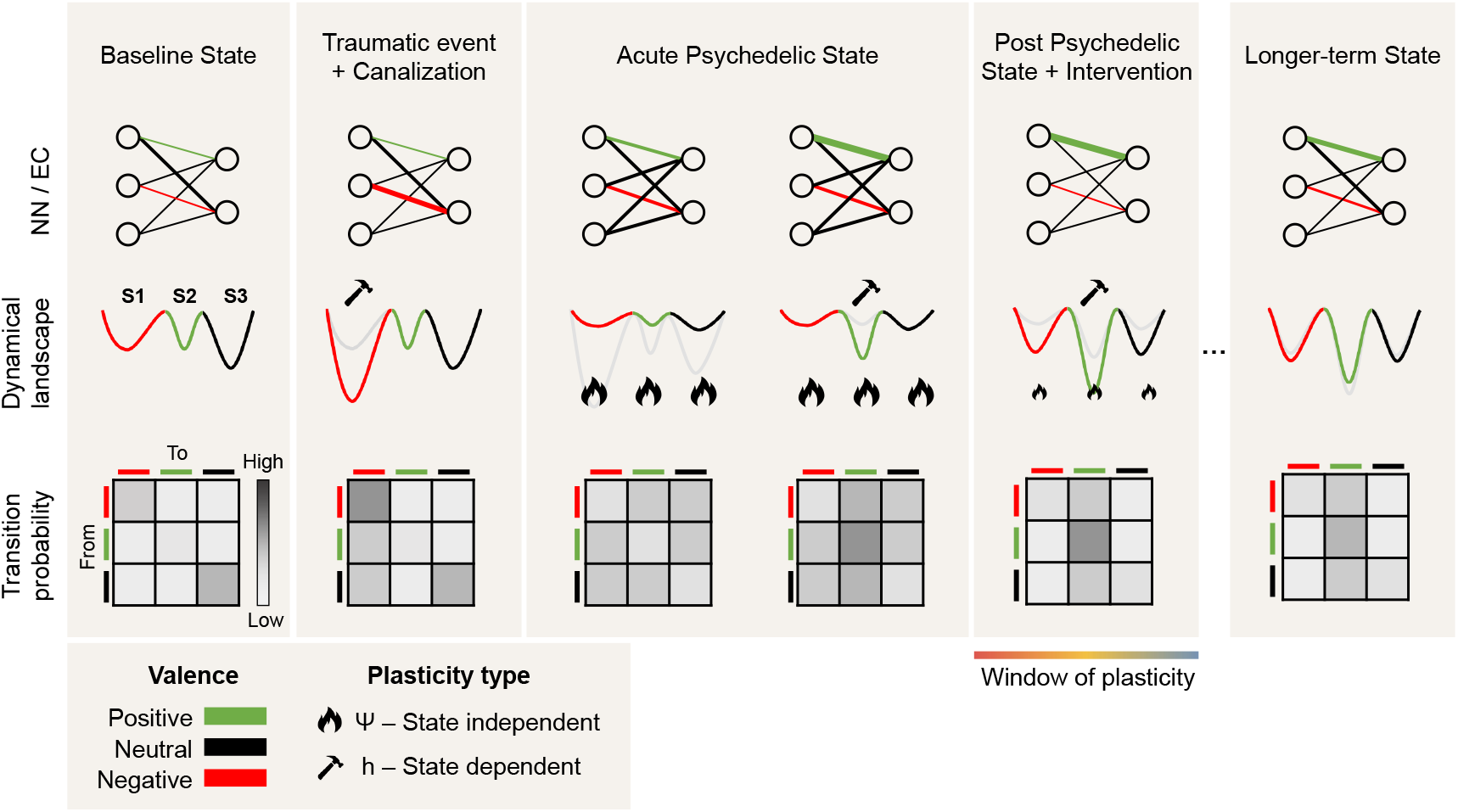
Psychedelics and psychopathology: a dynamical systems perspective. From left to right, we provide three views of the transition from health to canalization following a traumatic event and back to a healthy state following the acute effects and post-acute effects of psychedelics and psychotherapy. The top row provides the *neural network (NN) and effective connectivity (EC) view*. Circles represent nodes in the network and edge connectivity between them, with edge thickness representing the connectivity strength between nodes. The middle row provides *the landscape view*, with three schematic minima and colors depicting the valence of each corresponding state (positive, neutral, or negative). The bottom row represents the *transition probabilities across states* and how they change across the different phases. Due to traumatic events, excessive canalization may result in a pathological landscape reflected as a deepening of a negative valence minimum where the state may be trapped. During the acute psychedelic state, the landscape is deformed, enabling the state to escape. Moreover, plasticity is enhanced during the acute and post-acute phases, benefiting interventions such as psychotherapy or brain stimulation (i.e., changes in effective connectivity). Not shown is the possibility that a deeper transformation of the landscape may take place during the acute phase (see the discussion on the wormhole analogy in Section 4).

The interplay between external inputs, neural (fast time), and connectivity (slow time) dynamics can drive the system into a joint canalized, stable state of lower complexity. Under the influence of psychedelics, more diverse and complex dynamics destabilize the plasticity equilibrium point, leading to a more fluid and adaptable neural state in a process that is amplified by the plasticity-enhancing effects of psychedelics. This shift manifests as an acute systemic increase of disorder and possibly a longer-lasting increase in complexity (Ising temperature, Lempel-Ziv complexity, etc.) that affects both short-term dynamics and long-term plastic processes.

The CANAL framework offers insight into the neural mechanisms underlying the persistence of various brain disorders. In particular, psychedelics may mediate their effects by altering the balance between stability and plasticity in neural networks through meta-plasticity and thus act as potential therapeutic treatments. By acting on the serotonergic receptors, they trigger a cascade of neurochemical events, subsequently facilitating the reorganization of entrenched neural patterns. As discussed above, this alteration of the neural network during the acute phase (connectodynamics) can be interpreted as a rapid deformation or flattening of the landscape that allows the trapped state to escape and access more adaptive cognitive and emotional patterns. The rapid increase in complexity (a change in the dynamics) is in itself a likely driver of metaplasticity. The acute phase is believed to be followed by an extended window of malleability of the landscape, otherwise known as a “window of plasticity”, where treatments such as psychotherapy and transcranial electrical stimulation can further alter the pathological rigidity characteristic of various brain disorders (see Figure 4).

## 4 Neural geometrodynamics and general relativity

A parallel can be drawn between neural geometrodynamics and Einstein’s equations of general relativity — the original geometrodynamics. Both frameworks involve the dynamical interaction between structure and resulting activity, each influencing and being influenced by the other. The Einstein field equations, including the cosmological constant Λ, are

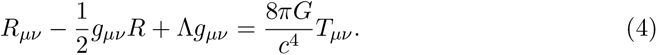

Here, *g*_*μ ν*_ is the metric tensor, *R*_*μ ν*_ = *R*_*μ ν*_[*g*_*μν*_] is the Ricci curvature tensor and a function of *g*_*μ ν*_, *R*[*g*_*μ ν*_ ] is the Ricci scalar (or curvature scalar) and a function of *g*_*μν*_, *T*_*μν*_ is the stress-energy tensor^3^, a function of the mass and energy distribution (all the indices refer to spacetime dimensions), *G* is the gravitational constant, *c* is the speed of light, and Λ is the cosmological constant. These equations describe the fundamental interaction of gravitation as a result of spacetime being curved by matter and energy. Specifically, they equate local spacetime curvature (on the left-hand side) with the local energy and momentum within that spacetime (on the right-hand side).

To complete these equations, the *geodesic equation* portrays how particles (matter) move in this curved spacetime, encapsulated by the notion that particles follow the straightest possible paths (geodesics) in curved spacetime,

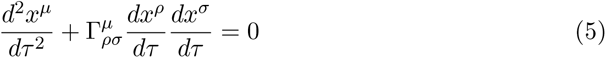

where *x*^*μ*^ are the coordinates of the particle, *τ* is the proper time along the particle’s path, and 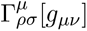 are the Christoffel symbols, which are a function of *g*_*μ ν*_ and encode the *connection* (a mathematical object that describes how vectors change as they are parallel transported along curves in spacetime). The stress-energy tensor *T*_*μ ν*_ can be computed from the state of the particles, closing the system of equations. For example, for *N* particles, it is given by 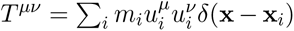, where *m*_*i*_ and *u*_*i*_ are the mass and velocity of the *i*th particle. More generally, the stress-energy tensor represents the state of matter and energy, which corresponds to *x* in our neural model. The metric *q*_*μν*_, which specifies the geometry of spacetime, is akin to the connectivity *w* — which shapes the structure of the space where fast dynamics occur.

In the context of neural mass models, the state equation, 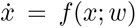, is analogous to the geodesic equation – “the state of the system evolves according to the landscape geometry specified by the parameters *w*”. On the other hand, the connectodynamics equation, 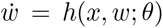 (with *θ* standing for plasticity parameters), is analogous to Einstein’s field equations — the parameters *w*, which describe the “structure” of the space where dynamics take place, evolve according to the current state of the system *x* and its ‘readiness’ for plasticity (parametrized by *θ*).

The analogy to psychedelic effects in general relativity can be clarified further. The neural effects of psychedelics, as we understand them, start with a disruption of connectivity in a spatially dependent manner. Since the analog of *w* is *g* (the metric), in cosmological terms, we would first see a dynamic deformation of spacetime independent of the mass distribution (state-independent plasticity). Spacetime would “flatten”. This would cause the mass in the universe to escape from gravitational wells following new geodesics (just as the state in the brain will explore new regions of phase space), in turn creating further deformations of spacetime (state-dependent plasticity).

We emphasize that this comparison is largely metaphorical and therefore limited: the mutual influence between particles and spacetime in general relativity is akin to the state of the neural system and its underlying connectivity parameters. In both cases, dynamics and structure are intertwined (see Figure 5). However, as an example of the limitations of the analogy, the slow and fast nature of the different variables is interchanged in the two formulations, with spacetime responding faster (at the speed of light) to changes in the distribution of energy than the stress-energy tensor itself.

**Figure 5:**
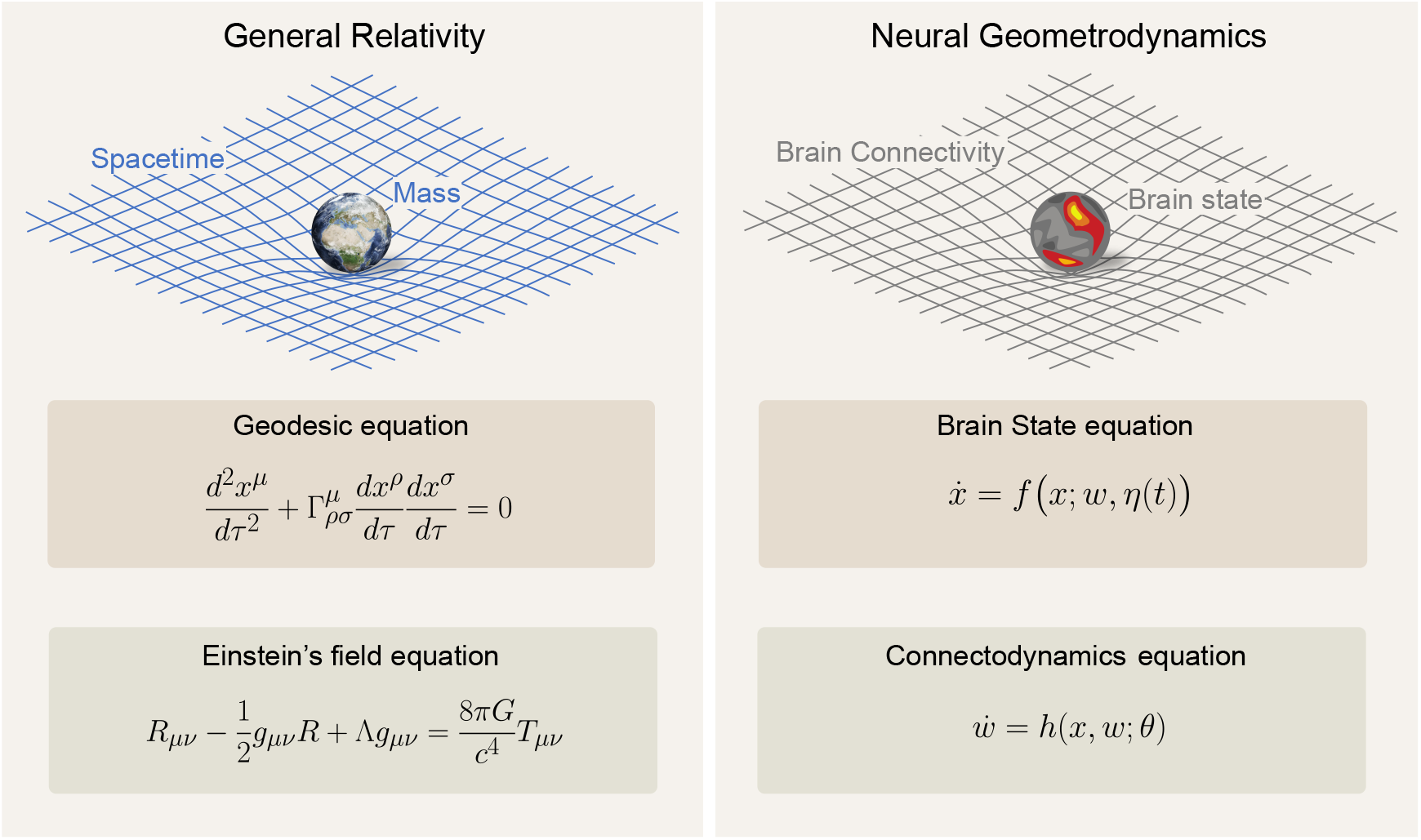
General Relativity and Neural Geometrodynamics. Left: Equations for general relativity (the original geometrodynamics), coupling dynamics of matter with the dynamics of spacetime. Right: Equations for neural geometrodynamics, coupling neural state, and connectivity. Only fast and slow time equations are shown (ultraslow time endows with dynamics the “constants” appearing in these equations).

### Metaplasticity and variable constants in cosmology

In our neural mass model framework, the concept of metaplasticity is introduced as dynamic variations in the plasticity control constants, namely *θ* in the connectodynamics equation. This set of constants can be represented as evolving over time as a function of the state of the system or other relevant variables,

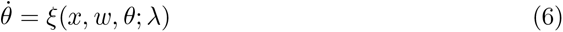

In this equation, *ξ* defines the evolution of the plasticity control constants with parameters *λ*.

Analogously, in the realm of general relativity and cosmology, it has been speculated that the fundamental constants, such as the speed of light *c*, the gravitational constant *G*, or the cosmological constant Λ, may, in fact, be dynamic. Although not part of the mainstream cosmological model, theories proposing variable constants, such as “Variable Speed of Light” (VSL) or “Variable Cosmological Constant” provide an intriguing parallel. For instance, within VSL theories, the speed of light *c* is postulated to vary over cosmological time scales. Certain hypothetical dynamical equations could dictate the dynamical evolution of these constants. Although these theories are quite speculative and do not form a part of mainstream physics, they offer an interesting perspective on the concept of metaplasticity and its potential implications for the dynamical evolution of neural mass models and the structure of their landscapes.

### Psychedelics as wormholes in the neural landscape

In the parallel of general relativity and neural geometrodynamics, we see the effects of psychedelics as a deformation of the neural landscape (spacetime) that allows the brain state (of a particle or set of particles) to escape from a local minimum and transition to another location in the landscape (spacetime). Although transitions may be smooth and respect the topology of the landscape (as described by topological quantities such as the Euler characteristic of Betti numbers^4^ [87, 88]), deformations of the landscape may also be more extreme — sharp transitions through a *topological tipping point* of the dynamical landscape. This may be due to external inputs (*η*(*t*)), when our system is non-autonomous [89], e.g., from sensory or brain stimulation effect. And as we have discussed, it may be due to connectivity dynamics.

The creation of a wormhole in general relativity^5^ can be viewed as a profound deformation of spacetime, bending and connecting distant parts of the universe in such a way that matter/energy, like an astronaut, can travel through vast distances in an instant. This change in the geometry and topology of spacetime can be likened to the effect of psychedelics on the human mind. Just as the wormhole alters the structure of spacetime, psychedelics may radically alter the dynamical landscape of neural dynamics, creating connections across distant landscape locations. In the same way that the astronaut uses the wormhole to bypass vast stretches of space, the deformation caused by psychedelics may allow the state of the brain to tunnel out and escape from a local minimum or stuck pattern of thought, providing access to new areas of the landscape — new perspectives and potentially unexplored territories of consciousness. This analogy, although speculative, aims to highlight that both phenomena are characterized by a fundamental transformation that enables traversal into otherwise inaccessible regions — whether in physical space or the brain’s dynamical landscape (see Figure 6 for a sketch of this concept).

**Figure 6:**
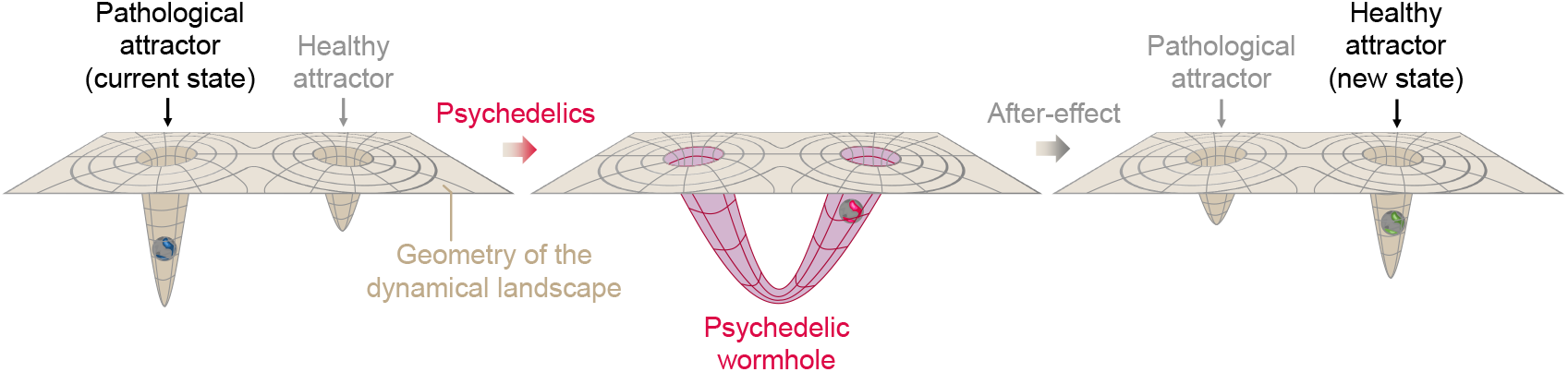
A hypothetical psychedelic wormhole. On the left, the landscape is characterized by a deep pathological attractor, where the neural state is trapped. After ingestion of psychedelics (middle), a radical transformation of the neural landscape takes place, with the formation of a wormhole connecting the pathological attractor to another, healthier attractor location and allowing the neural state to tunnel out. After the acute effects wear off (right panel), the landscape returns near its original topology and geometry, but activity-dependent plasticity reshapes it into a less pathological geometry.

### Characterizing the landscape

An important challenge in the program of neural geometrodynamics is to explore practical methods to characterize the landscape. Here again, we can draw inspiration from physics and mathematics.

The roots of this approach can be traced back to the 19th century when Carl Friedrich Gauss pioneered the field of differential geometry. Gauss’s Theorema Egregium demonstrated that the curvature of a surface could be determined entirely by measurements within the surface, without any reference to the surrounding space [92]. This seminal insight has laid the groundwork for understanding manifolds in various contexts, including the theory of relativity. In the era of general relativity, the interplay between geometry and physics was further enriched. Differential geometry and algebraic topology — which comes into play when one is interested in the global properties of the manifold, such as its shape, connectedness, and the presence of holes [93, 94] — became essential in describing the fabric of spacetime itself. It enabled physicists to conceptualize how mass and energy warp the geometry of spacetime, thus influencing the motion of objects.

In our current endeavor, these ideas find application in characterizing the complex dynamical landscapes of neural data. Modern tools from deep learning, such as variational autoencoders, can be used to unravel the reduced spaces underlying neuroimaging or neurophysiological data [53, 54], while dynamical systems theory in concert with differential geometry, group theory, and algebraic topology data analysis [95] offer robust frameworks to understand and characterize them [89, 96–100]. Topological data analysis can also be used to explore the graphs associated with model space, for example, the structural (connectome) or effective connectivity between regions in the brain (see [101] for a recent review). Topological methods have already been successfully employed to analyze detailed microscopic models [98], to study the relationship of criticality and topology in models [102], and to characterize functional brain networks derived from neuroimaging data [87, 88, 101].

World-tracking constraints force the brain as a dynamical system to mirror the symmetry in the data [37], a requirement that translates into constraints on structural and dynamical aspects of the system (and which can be analyzed using Lie group theory). This suggests leveraging the known links between topology and Lie groups [103]. The convergence of these mathematical techniques extends to neuroscience the fruitful exercise in physics of linking geometry and topology.

Finally, it would be interesting to explore if hierarchical data processing systems such as the brain display dynamical manifolds with hierarchical structure, including topology. This possibility is intuitive given the connections between the notions of criticality, information processing, and hierarchical organization [34, 104]. In this sense, the effects of psychedelics, which are seen to increase the temperature of the system [34] and the complexity of dynamics, should be reflected as an increase in the topological complexity of the associated dynamical attractors, as we discussed above with the analogy to wormholes.

The relationship between hierarchy and topological complexity could be analyzed, for example, by exploring artificial neural networks carrying out hierarchical processing (any generative deep network trained on real-world data would do, in principle). Such networks could then be used to generate neural activation data and analyze, for instance, whether the depth of the network (the number of layers in its hierarchical architecture) is reflected in the topology (e.g., in Betti numbers) associated with the data or its latent space.

## 5 Conclusions

In this paper, we have defined the umbrella of neural geometrodynamics to study the coupling of state dynamics and their complexity, geometry, and topology with plastic phenomena. We have enriched the discussion by framing it in the context of the acute and longer-lasting effects of psychedelics.

As a source of inspiration, we have established a parallel with other mathematical theories of nature, namely in general relativity, where dynamics and the “kinematic theatre” are intertwined (see the Appendix for a similar parallel with electrodynamics).

Although we can think of “geometry” in neural geometrodynamics as referring to the structure imposed by connectivity on state dynamics (paralleling the role of the metric in general relativity), it is more appropriate to think of it as the geometry of the reduced phase space (or invariant manifold) where state trajectories ultimately will lie: this is where the term reaches its fuller meaning. Since the fluid geometry and topology of the invariant manifolds underlying apparently complex neural dynamics may be strongly related to brain function and first-person (structured) experience [16], further research should focus on creating and characterizing these fascinating mathematical structures.

## Acknowledgments

GR, ELS, and RST have received funding from the European Research Council (ERC Synergy Galvani) under the European Union’s Horizon 2020 research and innovation programme (grant agreement No 855109). GR, ELS, JV, and RST have received funding from European Union’s Horizon 2020 research and innovation programme under grant agreement No 101017716 (Neurotwin).

## A Appendix

### A.1 A nautical analogy

To illustrate the interconnected dynamics of neural states, connectodynamics, and metaplasticity, consider a toy sailing boat navigating a circular pond. The boat moves through the pond, creating ripples that propagate across the water’s surface, eventually reflecting off the pond’s boundaries. These reflected ripples, in turn, influence the boat’s trajectory. This mirrors the dynamics of brain states, analogous to neural dynamics expressed by the equation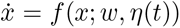, where the boat’s position represents the state *x* and the water surface’s geometry reflects the effective connectivity *w*. The term *η*(*t*) may be associated with an external force such as the wind.

The changes in the geometry of the water surface caused by the boat’s movement symbolize connectodynamics. This is captured by the plasticity equation 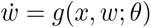, where the evolving connectivity parameters *w* depend on the boat’s position *x* and other factors. The boat’s position and the water’s surface geometry are intrinsically linked, akin to brain state and effective connectivity.

Further, imagine that other external factors, such as temperature fluctuations or changes in water viscosity, modify the water’s molecular structure over time. For example, a temperature decrease nearing freezing could alter the water structure (density and viscosity [105]) in the pond and how the boat’s movement affects the water geometry. This change in the water’s properties symbolizes the dynamics of plasticity, or metaplasticity, as described by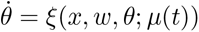.

### A.2 Classical dynamics of particles and fields

Here we provide the equations for other systems where one can think of part of the equation describing the geometry of a space-providing subsystem (“kinematic theatre”) and another the subsystem moving in this space, influenced by the structure and affecting its geometry in return. Several such examples can be found in physics.

#### Non-relativistic electrodynamics

The non-relativistic dynamics of *N* -charged particles and the associated electromagnetic field are described by (coupled) Newton’s second law and Maxwell’s equations. The charged particles are influenced by the electromagnetic field and at the same time, generate it. This can lead to problematic scenarios, such as the self-interaction problem: charged particles generate an electromagnetic field, and if one considers a particle’s interaction with its own field, paradoxical or unphysical results can arise. This self-interaction leads to divergences in calculations and has been a longstanding challenge in classical electrodynamics until recently [106].

The motion of the *i*th electron is given by 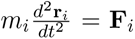, where **F**_*i*_ = *q*_*i*_(**E** + **v**_*i*_ *×* **B**) is the Lorentz force. The electromagnetic field obeys Maxwell’s equations:

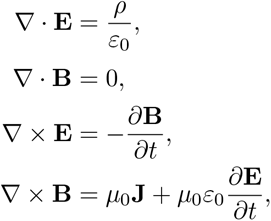

where *ρ*(**r**, *t*) = Σ _*i*_ *q*_*i*_ *δ* (**r** *−* **r**_*i*_(*t*)) and **J**(**r**, *t*) = Σ_*i*_ *q*_*i*_**v**_*i*_(*t*) *δ* (**r** *−* **r**_*i*_(*t*)) are the charge and current densities.

#### Relativistic equations

Maxwell’s equations are relativistic (they transform properly under Lorentz transformations), but Newton’s is not. For the **relativistic version**, the field strength tensor *F* ^*μ ν*^ is defined in terms of the four-potential *A*^*μ*^ [107],

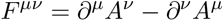

The dual tensor 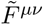 is defined in terms of *F*_*αβ*_ and the Levi-Civita symbol *ε*^*μ ν αβ*^,

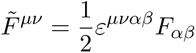

The homogeneous Maxwell’s equations are given by:

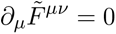

and the inhomogeneous Maxwell’s equations are given by:

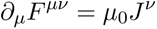

where *J*^*ν*^ is the four-current vector, which we now define for a particular case. Given a distribution of *N* particles each with charge *q*_*i*_ and four-velocity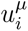, the four-current *J*^*μ*^ at position **x** and time *t* is given by

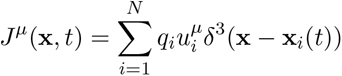

Here, **x**_*i*_(*t*) is the position of the *i*-th particle at time *t*, and *δ* ^3^ is the three-dimensional Dirac delta function.

The equation of motion for a charged particle in an electromagnetic field, commonly known as the Lorentz force equation, in its relativistic form is:

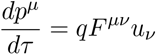

Here, *p*^*μ*^ is the four-momentum of the particle, *u*^*ν*^ is the four-velocity of the particle, *F* ^*μν*^ is the electromagnetic field tensor, and *q* is the charge of the particle. The equation describes how the four-momentum of the particle changes with proper time *τ* under the influence of the electromagnetic field.

### A.3 Modeling plasticity in neural mass models

In this section, we provide a brief overview of plasticity mechanisms and how they relate to the terms in the formalism, namely the functions *h* and *ψ*. See Table 1 for a summary. Including plasticity in Neuronal Mass Models (NMMs) allows for the modeling of time-varying connectivity strengths that reflect the learning and adaptation processes observed in biological neuronal networks.

**Table 1:** Summary of Different Types of Neural Plasticity Phenomena. **State-dependent Plasticity** (*h*) refers to changes in neural connections that depend on the current state or activity of the neurons involved. For example, functional plasticity often relies on specific patterns of neural activity to induce changes in synaptic strength. **State-independent Plasticity (***ψ*) refers to changes that are not directly dependent on the specific activity state of the neurons. For example, acute psychedelic-induced plasticity acts on the serotonergic neuroreceptors and thus acts on the brain networks regardless of specific activity patterns. Some forms of plasticity, like structural plasticity and metaplasticity, may exhibit characteristics of both state-dependent and state-independent plasticity, depending on the context and specific mechanisms involved. Finally, **metaplasticity** refers to the adaptability or dynamics of plasticity mechanisms.

| Type | Time Scale | Mechanism | Effects | State-dependence |
| --- | --- | --- | --- | --- |
| **Functional or dynamical Plasticity** | Milliseconds to minutes | Changes in strength/efficiency of synapses (e.g., Hebbian plasticity, LTP, LTD) | Short and long-term memory, fine-tuning of connections | State-dependent ($h$) |
| **Acute Psychedelic-induced Plasticity** | Minutes to Hours | Targeting of serotonergic neuroreceptors especially the 5-HT<sub>2A</sub> with a result in overall excitability [11, 12, 82, 83] | Flattens or de-weights the dynamical landscape | State-independent ($\psi$) |
| **Homeostatic Plasticity** | Hours to days | Regulation of overall excitability to maintain stability (e.g., adjusting synapse strength for E/I balance) [62–66] | Balances and stabilizes network | State-dependent ($h$) |
| **Structural Plasticity**<br>(e.g., post-acute psychedelic-induced plasticity) | Hours to years | Larger physical changes in neurons (e.g., dendrite growth, synapse formation) through an increase of endogenous BDNF and via TrkB binding (the receptor of BDNF) [84] | Long-term memory, development | Depends on context |
| **Metaplasticity** | Various, often longer-term | Changes in mechanisms governing synaptic plasticity (e.g., modulation of thresholds/rules) | Regulates other forms of plasticity, “plasticity of plasticity” [74] | Depends on context |

#### Functional Plasticity

The simplest and most common way to include synaptic plasticity is through Hebbian learning rules. Hebbian plasticity, a type of functional plasticity, follows the principle that “neurons that fire together wire together” [61]. It can be included in NMM using the equation

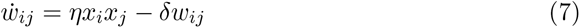

where *w*_*ij*_ is the synaptic strength from neuron *j* to neuron *i, x*_*i*_ and *x*_*j*_ are the neuronal activities, and *η* and *δ* are parameters controlling the learning and decay rates.

#### Homeostatic Plasticity

Homeostatic plasticity, a form of plasticity that adjusts synaptic strengths to keep the overall activity of a neuron or network within a certain range, can be included in an NMM using the equation [62–66],

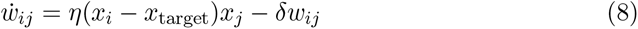

where *x*_target_ is the target activity level.

#### Structural Plasticity

Structural plasticity, where the actual number and dendrites and arrangement of synapses change over time, can be represented in NMMs by modifying (or even adding) rows and columns from the *w* adjacency matrix to represent the formation or elimination of synapses or even fibers.

#### Empirically-derived Structural Plasticity

NMMs can be used to infer the structural changes to plasticity without explicitly describing the plastic mechanism per se. For example, in the post-acute psychedelic state, NMMs can be used to infer the plastic changes to *w*_*ij*_ by optimizing the model functional connectivity 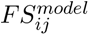 to approximate the empirical functional connectivity 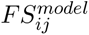 with a certain learning rate *ϵ* as in the following equation.

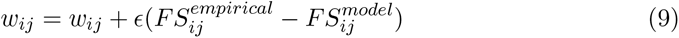

Such optimization is, for example, computed through gradient descent methods with priors on the topology of structural connectivity between brain regions [108]. Recent methods have further extended this framework by adding time-shifted correlation [42] to the optimization as a better description of the overall brain state, as in this case, the post-acute psychedelic state.

Including these forms of plasticity in NMMs allows for more realistic modeling of neural systems in better capturing their adaptive nature and the impact of learning and experience on synaptic connections.

**Figure A.1:**
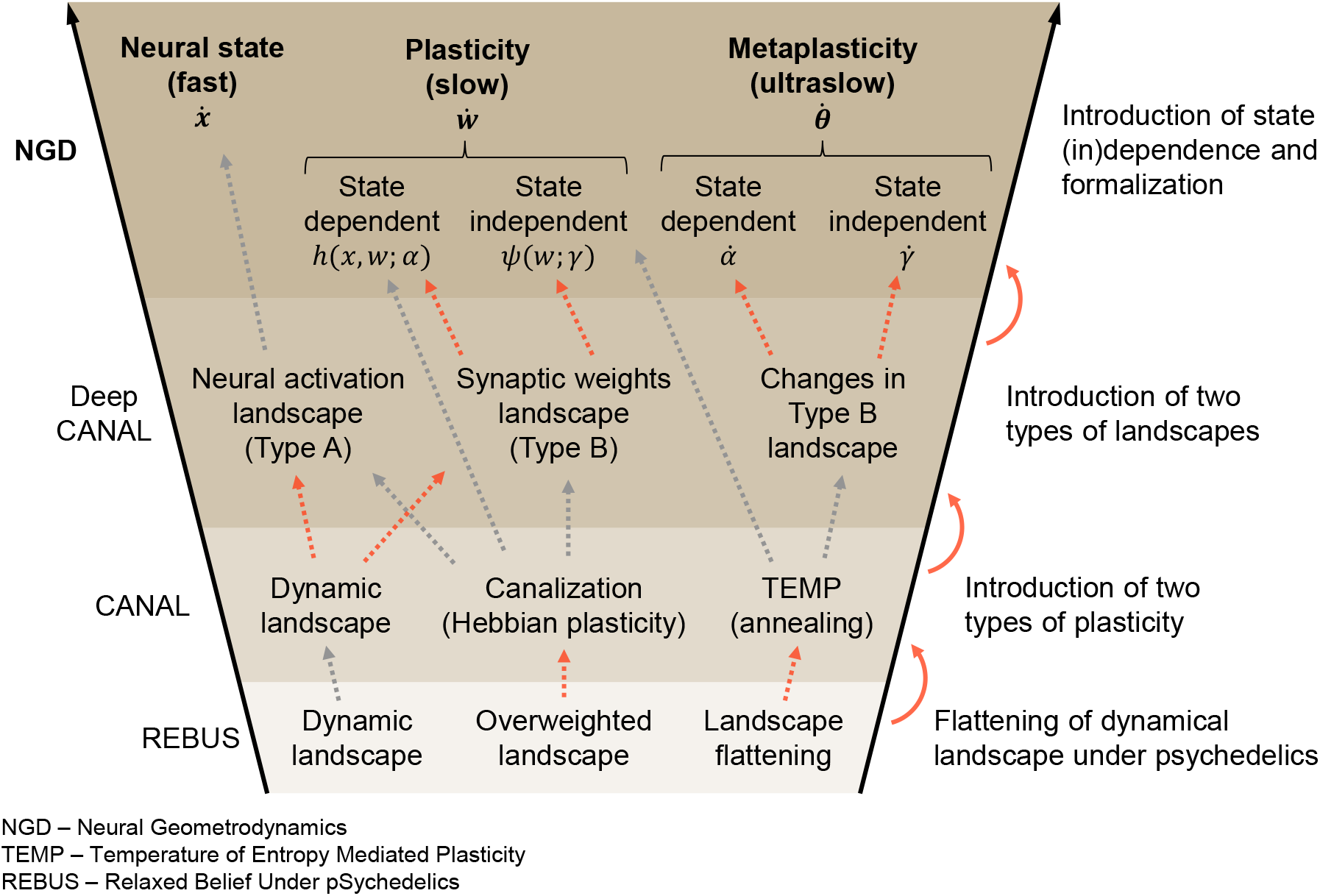
Conceptual funnel of terms between the NGD (neural geometrodynamics), Deep CANAL [48], CANAL [11], and REBUS [12] frameworks. The figure provides an overview of the different frameworks discussed in the paper and how the concepts in each relate to each other, including their chronological evolution. We wish to stress that there is no one-to-one mapping between the concepts as different frameworks build and expand on the previous work in a non-trivial way. In red, we highlight the main conceptual leaps between the frameworks. See the main text or the references for a definition of all the terms, variables, and acronyms used.

## Footnotes

1 In the REBUS model [56], from the Free Energy perspective, *w* would correspond to the weights or precision assigned to priors/beliefs; from the Entropic Brain perspective, *w* would correspond to the weights of the effective connectivity between neuronal populations on the macroscopic scale.

2 In the REBUS model and the Entropic Brain perspective [56], the weights of the effective connectivity during the psychedelic-induced state are “flattened” or “de-weighted”, representing a more symmetrical and non-hierarchical connectivity profile.

3 The stress-energy tensor (also called the energy-momentum tensor) is a central concept in general relativity. It encapsulates the distribution and flow of energy and momentum in spacetime, and its components include energy density, momentum density, and stress (pressure and shear stress) within a given region.

4 In algebraic topology, Betti numbers provide a way to count the number of *n*-dimensional “holes” in a manifold. The creation of a wormhole (in 4D or higher dimensional spaces), being a topological feature that connects two otherwise distant regions of spacetime, would alter the topological structure of the manifold it inhabits and the associated Betti numbers.

5 Wormholes, a term due to John A. Wheeler [90], also known as Einstein-Rosen bridges, are solutions to the Einstein field equations of general relativity which some models suggest could exist under certain conditions. However, creating or stabilizing a traversable wormhole would likely require forms of exotic matter with properties not yet observed in the known universe — there is no current consensus about this in classical general relativity, where some theorems suggest it may not be possible in some conditions because of the necessity of singularities, or in quantum gravity, where topology change is a natural concept [91].

